# A Self-Supervised Workflow for Particle Picking in Cryo-EM

**DOI:** 10.1101/2020.03.13.991471

**Authors:** Donal M. McSweeney, Sean M. McSweeney, Qun Liu

## Abstract

High-resolution single-particle cryo-EM data analysis relies on accurate particle picking. To facilitate the particle picking process, we have developed a self-supervised workflow. Our workflow includes an iterative strategy to use the 2D class average to improve training particles and a progressively improved convolutional neural network (CNN) for particle picking. To automate the selection of particles, we define a threshold (%/Res) using the ratio of percentage class distribution and resolution as a cutoff. Our workflow has been tested using six publicly available data sets with different particle sizes and shapes, and is able to automatically pick particles with minimal user input. The picked particles support high-resolution reconstructions at 3.0 Å or better. Our workflow offers a way toward automated single-particle Cryo-EM data analysis at the stage of particle picking. The workflow may be used in conjunction with commonly used single-particle analysis packages such as Relion, cryoSPARC, cisTEM, SPHIRE, and EMAN2.

## 1. Introduction

The rapid development of computational algorithms and workflows has boosted the resolution revolution in high-resolution single-particle cryo-electron microscopy analysis (Cheng, 2015; Henderson, 2015; Subramaniam *et al.*, 2016). With the further improvement on electron microscope optics, camera speed, and data collection strategies, collecting 4000-10000 micrographs per day is becoming routine. Of course this improvement has resulted in substantial amounts of data to be processed, and it becomes time consuming to go through each of these steps in single-particle analysis workflows implemented in program packages such as Relion, cryoSPARC, cisTEM, SPHIRE, and EMAN2 (Tang *et al.*, 2007; Fernandez-Leiro & Scheres, 2017; Moriya *et al.*, 2017; Punjani *et al.*, 2017; Grant *et al.*, 2018). These packages have a particle picking process either manually or semi-automatically. However, finding suitable parameters for automated particle picking remains difficult, a situation which is amplified when dealing with low contrast micrographs with contamination or denatured particles. Traditional methods involve manually picking particles and using manually selected 2D class averages in order to obtain accurate templates for template-based automated particle picking (Frank & Wagenknecht, 1984; Huang & Penczek, 2004; Chen & Grigorieff, 2007; Tang *et al.*, 2007; Langlois *et al.*, 2014; Scheres, 2015; Punjani *et al.*, 2017). Each of these steps may require expert knowledge to judge the quality of particles and to choose, on a trial-and-error basis, parameters for template-based particle picking. In an additional complication, with low contrast micrographs such as close-to-focus ones that preserve high-resolution information, picking particles manually can be non-trivial and laborious even for experts.

Due to rapid accumulation of large cryo-EM data sets, using automated particle picking to facilitate single-particle analysis is highly desirable (Danev et al., 2019). Convolutional neural networks (CNN) have been increasingly used for particle picking in cryo-EM single-particle analysis (Wang *et al.*, 2016; Xiao & Yang, 2017; Zhu *et al.*, 2017; Bepler *et al.*, 2018; *Da et al., 2018*; Nguyen *et al.*, 2018; Al-Azzawi *et al.*, 2019b; Wagner *et al.*, 2019). These CNN-based methods may differ in the formation of network architecture. Nevertheless, they all require particle data for training and the training quality determines the picking results and subsequent single-particle analysis. The training data can be composed of either manually picked particles or *ab initio* picking by various feature detection methods (Zhu *et al.*, 2004; Voss *et al.*, 2009; Al-Azzawi *et al.*, 2019a). However, even for these methods that claim to be automated, they require using pre-trained CNN models. These models may not be always reliable for unknown particles due to data set bias (Wang *et al.*, 2016; Tegunov & Cramer, 2019; Wagner *et al.*, 2019).

An effective strategy is needed such that CNNs can be trained in a self-supervised manner for improved particle picking. Considering the established utility of 2D class averages in selecting particles and CNNs in pattern recognition, we propose that the combination of the two could improve the quality of training data via iterative training, particle picking, and 2D class averaging. To test this hypothesis, we devised a self-supervised iterative particle picking workflow that may be used for automated particle picking and can be incorporated into a variety of single-particle analysis packages. Here we describe the process and performance of the workflow, which we have tested with six data sets that span a variety of particle sizes and shapes. We offer some ideas for further enhancement of the use of our workflow.

## 2. Methods

### 2.1 Cryo-EM micrograph data preparation

We used six publicly available EMPIAR (https://www.ebi.ac.uk/pdbe/emdb/empiar/) data sets to test the workflow as summarized in **Table 1**. Among these data sets, EMPIAR 10204, 10218, 10028, and 10335 are unaligned movies. We used 5 by 5 patches and reported dose rates for dose-weighted motion correction in Relion (Zivanov et al., 2018). Data sets EMPIAR 10184 and 10059 were already motion corrected and were used directly for downstream use. Per-micrograph CTF correction for both phases and amplitudes were performed in Gctf (Marabini et al., 2015; Zhang, 2016). After CTF correction, we selected aligned micrographs with an estimated CTF resolution beyond 3.0 Å for EMPIAR 10204 and 4.0 Å for the others to test our workflow. To generate a subset of micrographs for iterative training and particle picking, we selected 20-40 micrographs half of which had defocus below 1 μm and the others had defocus below 2 μm. For EMPIAR 10204, we used the first 20 micrographs to test our workflow.

**Table 1.** Summary of results for EMPIAR data sets used in this work.

| EMPIAR | EMD | Name | MW (kDa) | # Particles picked | # Particles refinement | Percent (%)<sup>§</sup> | Resolution (Å) | B-factor (Å<sup>2</sup>) |
| --- | --- | --- | --- | --- | --- | --- | --- | --- |
| 10204 | 6840 | β-galactosidase | 520 | 58710 | 31542 (93975) | 53.7 | 2.66 (2.6) | 65 |
| 10218 | 9233 | 20S proteasome | 700 | 80346 | 49870 (127570) | 62.1 | 2.4 (2.1) | 68 |
| 10184 | 7550 | Aldolase | 150 | 922306 | 114133 (187000) | 12.4 | 2.45 (2.4) | 107 |
| 10059 | 8117 | TRPV1 with DkTx and RTX | 280 | 441246 | 91651 (73929) | 20.8 | 3.0 (2.95) | 113 |
| 10028 | 2660 | 80S ribosome | 1263 | 155264 | 118801 (105247) | 76.5 | 2.85 (3.2) | 98 |
| 10335 | 20907 | Streptavidin | 53 | 691567 | 12206 (11402) | 1.8 | 2.69 (2.6) | 58 |

In parentheses are reported values in the database.
<sup>§</sup>Percentage ratio of the number of particles used in the final refinement and the number of particles picked.

### 2.2. A workflow for iterative particle picking

The workflow is built on the hypothesis that from a subset of micrographs, particles may be improved by selectively filtered through 2D class average and the improved particles are then used to train a CNN network. We propose that this iterative procedure will lead to the optimization of a fine-tuned CNN-based particle picker, capable of picking high-quality particles. The workflow is composed of three steps as illustrated in **Fig. 1**. The first step produces initial candidate particles for training the CNN network. The second step trains the network progressively, leading to the final particle picking in step 3. To speed up the convergence of the CNN model, 2D class averages are used to produce improved particles (**Fig. 1a**). The selection of 2D classes and particles is automated by using the ratio of percentage class distribution and resolution (denoted as %/Res), both of these statistics are reported in the Relion model star file. These particles in the selected 2D classes were then used for iterative 2D class averaging and selection until 90% of particles are selected (**Fig. 1a**).

**Figure 1.**
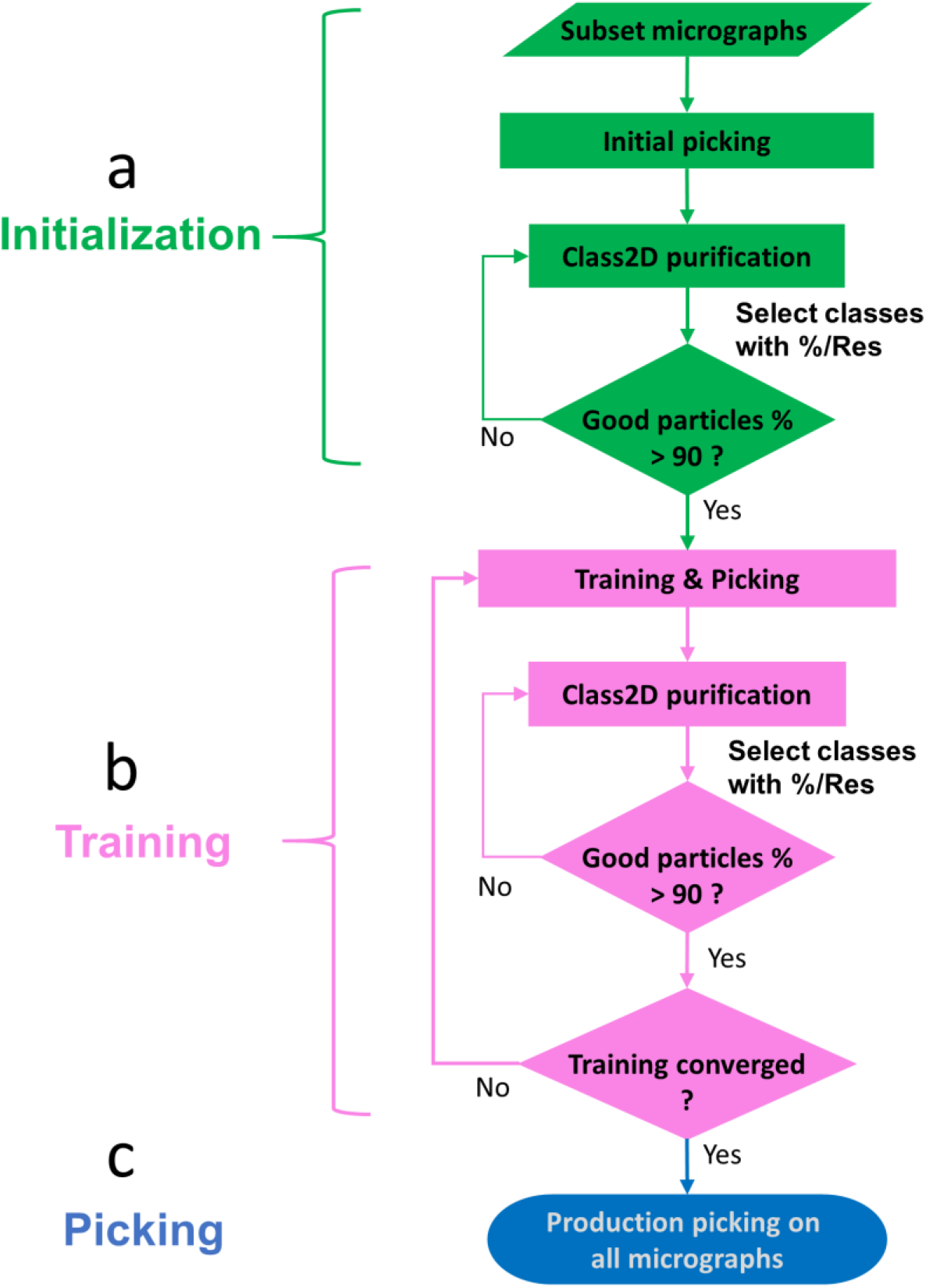
Schematic drawing for the workflow of iterative particle training and picking. A small set of data, usually 20-40 micrographs, is used for the workflow. The workflow is comprised of an initialization step (**a**), a training step (**b**) followed by a picking step (**c**). The process uses iterations of 2D class averages to improve particles for training a convolutional neural network (CNN) for particle picking.

Even with the use of 2D class averages, these initial particles may not be optimal chosen which may lead to subsequent biased training and picking. Therefore, following training and picking, we perform 2D class averaging again to improve particles until 90% of particles exceed the %/Res threshold. The training, picking, and 2D class average are repeated until convergence. Here, the definition of convergence is based on the ratio of the number of qualified particles (i.e. exceeded %/Res cutoff) to the total number of picked particles. In this work, we used 70% as a termination cutoff for convergence. That is, if after training, 70% of picked particles are in selected 2D classes with %/Res > 0.1, we consider the training converged and the trained network is then used for a production picking.

### 2.3 *ab initio* particle picking

To produce candidate particles for training, we implemented a Localpicker for *ab initio* particle picking. The program makes use of a threshold mask image that is calculated based on the value of local pixels (Singh et al., 2012). With the threshold mask image, features were detected, labeled, and written to a star file, one file for each micrograph. One particular feature of Localpicker is that it is a shape-based method, thus enabling the picking of particles of various shapes simultaneously. Localpicker is robust and requires only three parameters to control the particle picking process: estimated particle size in pixels, bin size, and threshold. The particle size is used to remove particles that are too close on micrographs. The bin size is used to reduce micrograph size to facilitate the picking. The threshold is used for feature detection; and local maxima smaller than the threshold value are ignored.

For five EMPIAR data sets 10204, 10184, 10059, 10028, and 10335, we used Localpicker for initial particle picking with bin size 9 and threshold 0.001 or 0.0015. For EMPIAR 10218 (20S proteasome), due to aggregation among particles, we manually picked ~1000 particles for downstream workflow.

### 2.4 Initial particle selection

Initial particles picked manually or by Localpicker were extracted from micrographs and scaled to 64×64 pixels using Relion (Version 3.0.7) followed by iterative 2D class averaging and selection of 2D classes. The number of classes used for 2D classification is the total number of particles divided by 200. The selection of 2D classes was based on %/Res. Only these classes with a %/Res value of greater than 0.1 were selected for the next cycle of 2D class average. The 2D class averaging and particle selection were iterated until more than 90% of particles were selected (i.e more than 90% of picked particles reach the aforementioned cutoff value).

### 2.5 CNN architecture

For our particle picking workflow, we employed a three-convolution-layer network architecture (**Fig. 2a**). The network contains an input layer, three layers of convolution (Conv2D) followed by a pooling operation (MaxPooling2D) for feature extraction at various scales. Finally, two densely connected layers are used for input classification. The last dense layer has two outputs, whose values correspond to the relative probability of classification as a particle or a non-particle. Given a candidate image, the network assigns a probability of being a particle and non-particle with a summed probability of 1.

**Figure 2.**
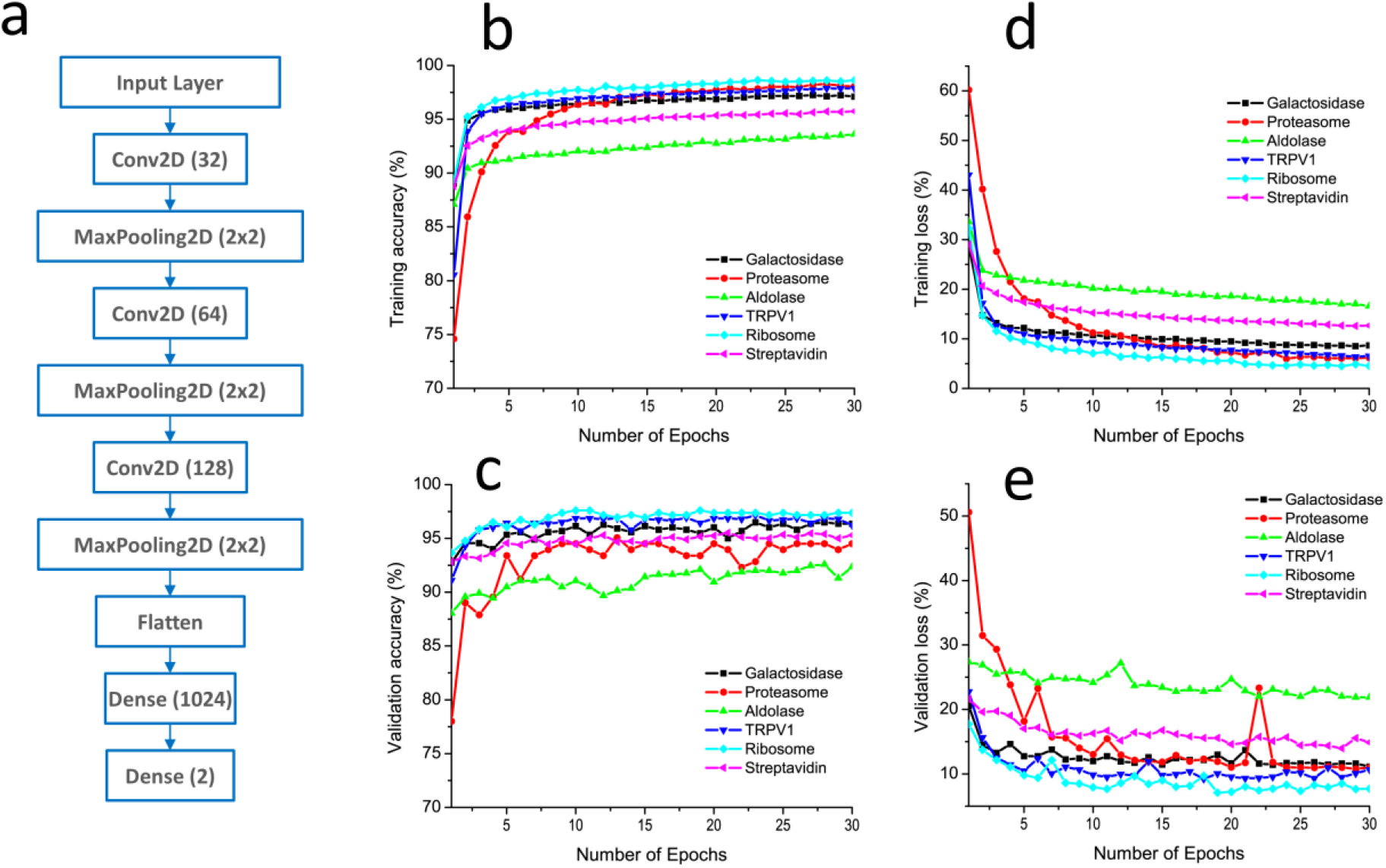
CNN network for training and picking. (**a**) A three-convolutional-layer CNN architecture. The three Conv2D layers use indicated number of filters in parentheses. After each Conv2D, the spatial dimensions of the filters are reduced by a factor of 2 through the pooling process. Two Dense layers are used to classify candidate particles. (**b-e**) Training results for the six EMPIAR data sets demonstrate the convergence of the workflow. (**b**) Training accuracy. (**c**) Validation accuracy. (**d**) Training loss function. (**e**) Validation loss function.

### 2.6 Iterative training and particle picking

Selected particles from the initialization stage were used for training the CNN implemented using Keras (https://keras.io) with TensorFlow as the backend. For the training and picking, we binned particles by 4 and resized them to 64×64 pixels. For each iteration, the training was performed for 30 epochs and the training accuracy, validation accuracy, training loss and validation loss were monitored for convergence. No parameters were specially tuned during the iterative training and particle picking processes. We coded the Keras-based particle training and picking as a program named as Kpicker.

Data augmentation was used to synthesize additional data to facilitate the training. Specifically, we used random rotation of 20 degrees and flips (vertical and horizontal) for augmentation. Particles selected from 2D class average were labeled as 1. Non-particles were randomly selected from empty areas at a minimum distance of a particle diameter from known particles. These non-particles were labeled as 0.

To predict whether a candidate image is a particle or not, we optimized the model with respect to the binary cross-entropy loss where a softmax activation function was used on the final layer. Kpicker scans over micrographs to produce a stack of candidate images of 64×64 pixels in size. These images were provided to the model to get predications as particles or non-particles. We treat a candidate image as a particle if its binary classification probability is 0.9 or higher. When the two particles are too close to each other, we keep the particle with a higher predicted probability. The same 2D class average was used to filter particles and %/Res of 0.1 was used for automatic selection of 2D classes. In general, two iterations of Kpicker training and picking followed by 2D class averaging are sufficient for reaching convergence. The CNN was then used for production picking against all micrographs. **Table 1** summarizes the number of particles picked for the six test data sets.

### 2.7 Reconstruction of 3D maps

Picked particles were extracted as 64×64 pixels and further cleaned up by 2D class averages in Relion (Zivanov et al., 2018) or cryoSPARC (Punjani et al., 2017). Cleaned-up particles were re-centered and re-extracted with a bin-size 2 for EMPIAR 10335 and 1 for the other data sets. These particles were used for 3D classifications and high-resolution refinements. Appropriate symmetry was enforced for all refinements except the *ab initio* 3D reconstructions in which none symmetry (C1) was used. The local map resolution was estimated using ResMap (Kucukelbir et al., 2014). B factors of the reconstructed maps were estimated using a Guinier plot (Rosenthal & Henderson, 2003).

## 3. Results

### 3.1 Training and picking with the workflow

Kpicker in our workflow contains a training and a picking module. To speed up the training process, we down-sized all particles to 64×64 pixels for all six data sets. Particles from these subset micrographs were used for training the network for production picking. Within 30 epochs, the training process had converged signaled by a plateau in both the accuracy and loss (**Figs. 2b-2e**). With the filtered particles from the iterative 2D class averages, the training process is quite robust with an accuracy beyond 0.9 (**Figs. 2b & 2c**). Among these six test data sets, ribosome data show the best validation performance (accuracy and loss) while aldolase and streptavidin data lead to relatively poorer performance. Considering that the mass of the ribosome is 1.3 MDa; and streptavidin and aldolase have masses below 200 kDa, such divergent performance might suggest a particle-size dependent training efficiency. This is consistent with the fact that large particles have higher signal-to-noise ratios compared to particles of smaller sizes.

To visualize the picking quality of our workflow, we show two representative micrographs with picked particles for the β-galactosidase data (EMPIAR 10204) (**Fig. 3**). The two micrographs contain some ice contaminations. With a particle size of 240 pixels in diameter (212 Å), a bin size 9, and a threshold 0.0015, Localpicker effectively picked most particles. However, these ice contaminants were also picked due to their high intensities. After the iterative training and picking, Kpicker can effectively classify these ice-contaminated areas as non-particles, leading to improved picking. Such improved picking capability is likely due to the improved training data and hence improved CNN model.

**Figure 3.**
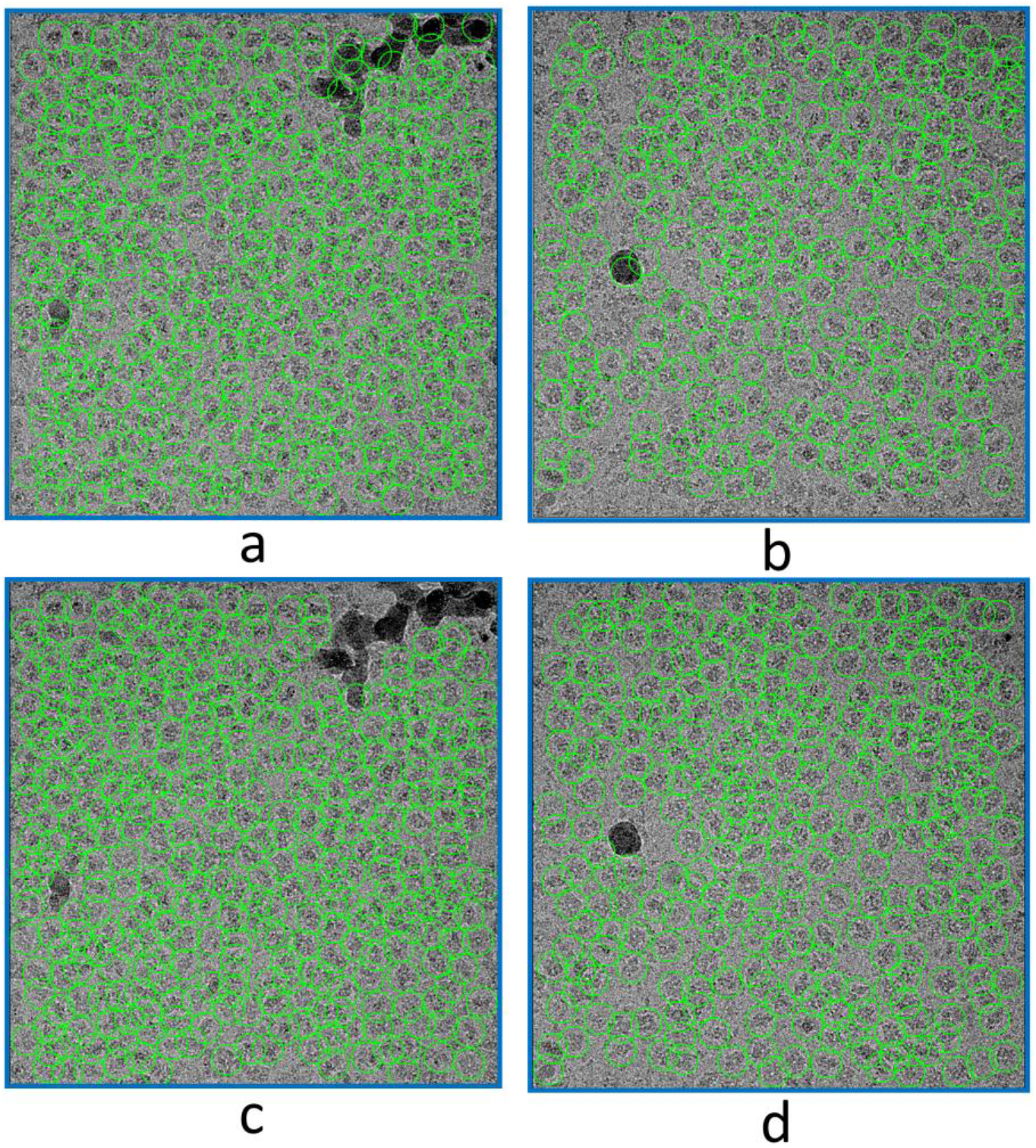
Particle picking before and after iterative training and picking. Picked particles are indicated as green circles. (**a, b**) Two representative micrographs with particles picked initially by Localpicker. (**c, d**) Improved particle picking after the iterative procedure of the workflow. With the workflow, iced areas have been effectively excluded from picking.

It is noted that, with the six data sets, we did not adjust training and picking parameters used for Kpicker except particle size. For each data set, the particle size used in Kpicker was the same as that used for Localpicker or manual picking. Therefore, our workflow and the CNN network are promising for allowing self-supervised training and picking across multiple data sets with minimal required adjustments to support single-particle cryo-EM data analysis.

### 3.2 Use of %/Res criterion for automated 2D class selection

In our workflow, an important step is the selection of particles from 2D classes for subsequent training. In general, good classes have a higher percentage class distribution and a higher resolution (a smaller value). Instead of using a single criterion, either class distribution or resolution, we choose to use their ratio (%/Res) to filter 2D classes. **Figure 4** shows the distribution of %/Res with respect to the number of classes for the six test data sets. We found that %/Res gives a sharp contrast between the number of good and bad classes and can be used to select classes and thus particles automatically. For five data sets, %/Res decreases rapidly before reaching a value of 0.1. The only outlier is the streptavidin data in which more than 50 classes have a %/Res of over 0.1. Tetrameric streptavidin is a small protein of 53 kDa. A wider %/Res distribution is consistent with a lower accuracy in alignment of particles within each 2D class. Nevertheless, we found a %/Res of 0.1 is a good compromise for selecting promising 2D classes automatically, including streptavidin, for training the network to convergence (**Figs. 2b-e**).

**Figure 4.**
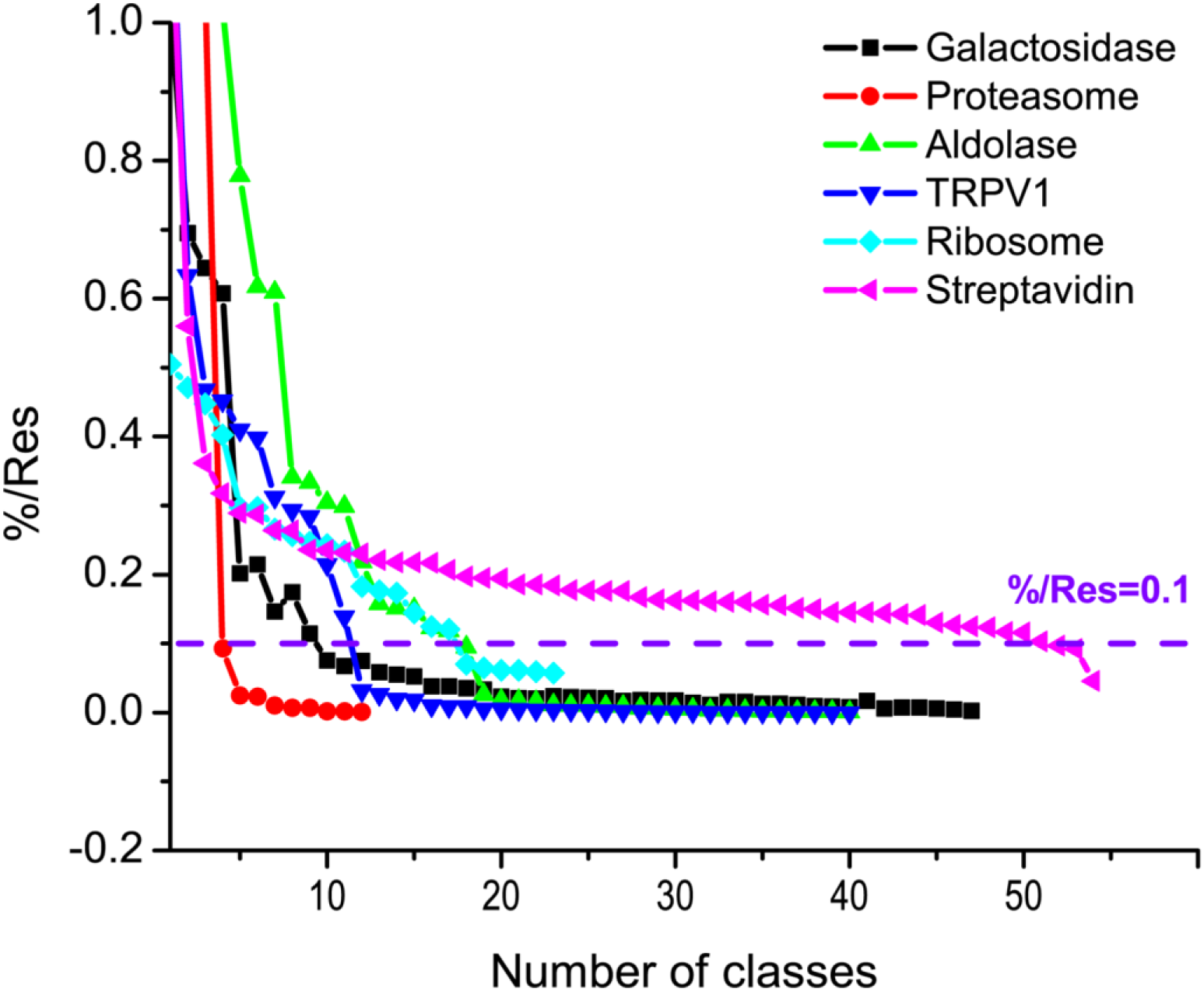
Plot of %/Res with respect to the number of classes for the six EMPIAR data sets. Particles iteratively trained by the CNN model and improved by 2D class average until 90% or more particles were selected by %/Res of 0.1 (dashed purple line).

With the established workflow and the automated 2D class selection criterion of 0.1 %/Res, we performed the production picking in Kpicker for all micrographs for the six data sets, each with an individually trained network and a respective particle size. **Table 1** summarizes the total number of particles picked for each data set, ranging from 59 K particles for β-galactosidase (213 micrographs) to 922 K particles for aldolase (1100 micrographs).

### 3.3 High-resolution 3D reconstructions

To test whether our workflow and picked particles support high-resolution single-particle analysis. We performed 2D and 3D classifications and 3D refinements for the six data sets from the picked particles. With particles extracted and scaled to 64×64 pixels, 2D classification reveals clear classes with distinctive molecular shapes and atomic features for all six data sets (**Fig. 5**). For TRPV1 channels in protein nanodiscs, the contrast between the channels and the disks allow the appreciation of the embedded transmembrane regions (**Fig. 5e**). At this 2D classification stage, we selected particles with distinctive 2D features for 3D classifications and refinements.

**Figure 5.**
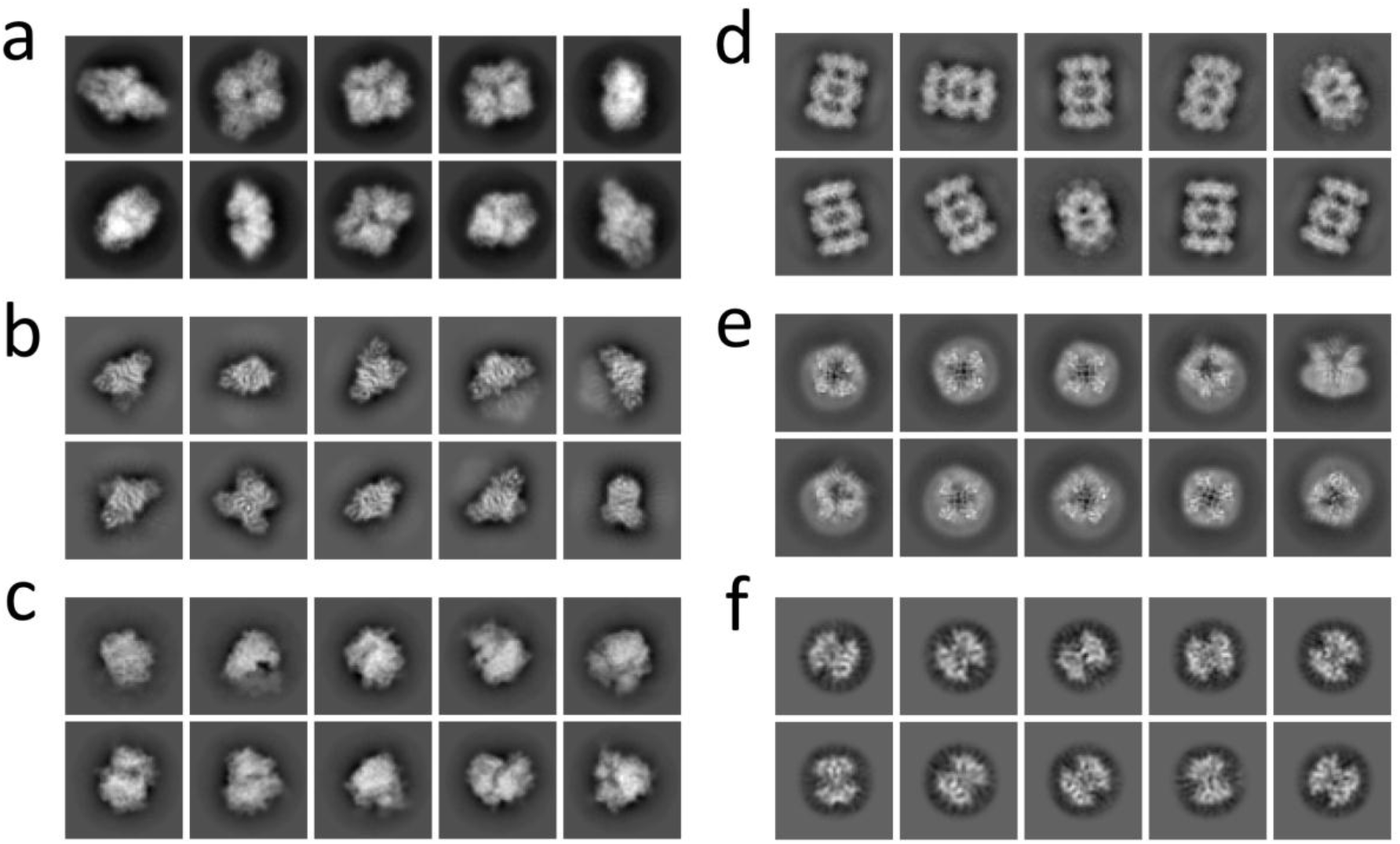
Representative 2D classes for the six EMPIAR data sets. Particles picked by the workflow were filtered by 2D class averages. (**a**) β-galactosidase. (**b**) 20S proteasome. (**c**) Aldolase. (**d**) TRPV1. (**e**) 80S ribosome. (**f**) Streptavidin.

To further test whether picked particles support high-resolution reconstructions, we re-centered and re-extracted these selected particles from micrographs and performed 3D classifications and refinements for achieving a high resolution. Using a gold standard Fourier shell correlation at 0.143 as a cutoff, particles from all six test data sets are readily refined to maps of a resolution of 3 Å or better: 3.0 Å for TRPV1 embedded in protein nanodiscs and 2.4 Å for proteasome (**Fig. 6**). The number of particles used for their final refinements is listed in **Table 1**. Their local resolution maps indicate high-resolution features (**Figs. 7a-f**). These test data sets cover diverse samples of different shapes and sizes from 1.3 MDa ribosome to 53 kDa streptavidin. Compared to the reported resolutions in the database, particles from our workflow allowed the 3D reconstructions at equivalent resolutions (**Table 1**).

**Figure 6.**
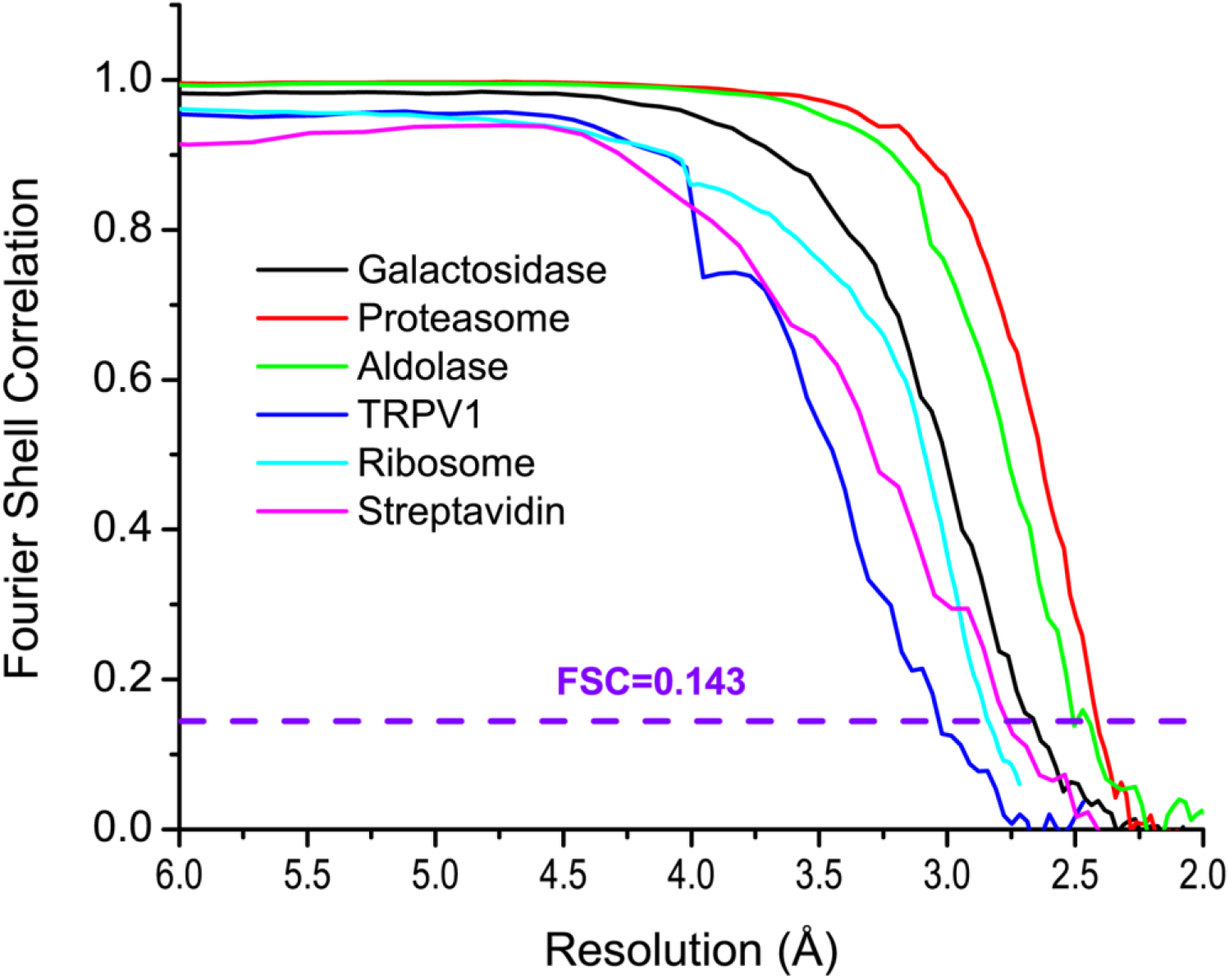
Gold-standard Fourier shell correlation for the six EMPIAR data sets. The dashed purple line indicates the cutoff of FSC at 0.143.

**Figure 7.**
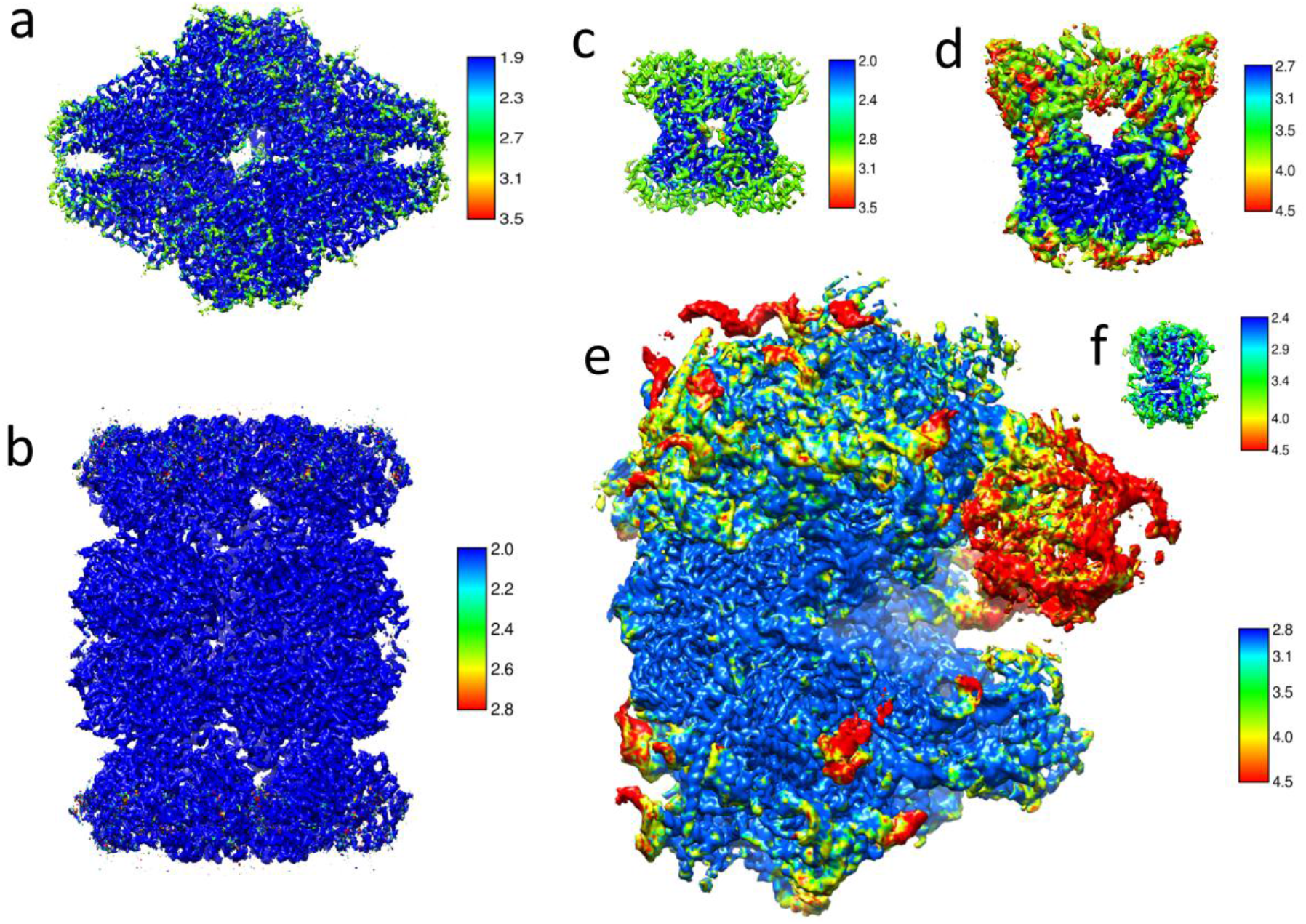
Refined 3D maps for the six EMPIAR data sets. For each data set, refined maps were color coded based on local-resolution estimation. The size of each reconstruction roughly reflects its actual size relative to ribosome. (**a**) β-galactosidase (520 kDa). (**b**) 20S proteasome (700 kDa). (**c**) Aldolase (150 kDa). (**d**) TRPV1 (280 kDa). (**e**) 80S ribosome (1263 kDa). (**f**) Streptavidin (53 kDa). The keys indicate the resolutions of the colored maps: darker blue for higher resolution; and darker red for lower resolution.

To evaluate our workflow relative to other particle picking programs such as Relion, we used the deposited particles of the ribosome data (EMPIAR 10028). These ribosome particles (105247) had been picked using Relion and were refined to 3.2 Å resolution (Wong *et al.*, 2014). With our workflow, Kpicker picked 155264 particles from the ribosome data. If we take these shiny particles as ground truth, 95.8% of them (100841) were picked by Kpicker with their coordinate centers within 40 pixels. From these Kpicker picked particles, a 2.84 Å reconstruction can be readily obtained (**Table 1**).

Therefore, our workflow, including the use of 0.1 %/Res selection criterion, can pick particles of sufficient quantities of high-quality particles to support high-resolution Cryo-EM data analysis.

## 4. Discussion

### 4.1 Particle picking

In this work, we proposed and tested a deep learning based iterative workflow to facilitate particle picking and improvement for Cryo-EM single-particle analysis. With a prior knowledge of particle size (in pixels), the particle picking process can be automated from initial particle selection to filtering by 2D class averages and finally to large-scale production picking (**Fig. 1**).

In our workflow, we used Localpicker and 2D class average to generate initial particles for Kpicker training. One can also pick particles manually, as was done for proteasome data (EMPIAR 10218), and use them for Kpicker training with or without 2D class average.

There are no limitations on the number of particles to be used for training. With the β-galactosidase data as an example, 100 particles give decent training and picking results although more particles are beneficial. We found that 50 particles can lead to the picking of 13% of particles with their coordinate centers within 20 pixels of finally refined ones. This number increased significantly to 65% when 100 particles were used for training. Therefore, we suggest using at least 100 particles for Kpicker training. With the iterative 2D class average and training procedure, Kpicker tends to pick more and improved particles. For the production picking of the β-galactosidase data with 4656 training particles, 81% of particles have their coordinate centers within 20 pixels of the final refined positions.

We have developed two pickers, Localpicker and Kpicker, for testing with our workflow. Both pickers take MRC format micrographs and write out particle coordinates in star format. Therefore, they may be used alone to pick particles for other workflows. In the current implementation, the two pickers have their limitations. For example, we haven’t implemented an ice detection step. Therefore, ice contaminated area may be picked by Localpicker. Nevertheless, these false particles were rejected from 2D class average and skipped by Kpicker (**Fig. 3**). The effectiveness of excluding ice area from Kpicker indicates the utility of our workflow in facilitating single-particle analysis. One can also include ice areas as negative particles for training as used by FastParticlePicker (Xiao & Yang, 2017). As a proof of concept, the current version of Kpicker training makes use of GPU while the picking is CPU only. To speed up the Kpicker picking performance, picking in GPU is more desirable (Wagner et al., 2019).

For the six test data sets of various pixel sizes ranging from 0.536 Å (streptavidin) to 1.34 Å (ribosome), we found that a bin size between 7-9 and a threshold of 0.001-0.0015 yield decent picking results by Localpicker. If one needs to optimize the initial picking, altering the bin size and threshold is recommended. In Localpicker, there is almost no need to change the particle size because it is only used for cleaning up close-contract particles and does not contribute to pattern recognition.

### 4.2 Low-defocus micrographs

For the β-galactosidase data, we selected the first 20 micrographs for iterative training and picking. We found that for low defocused micrographs (defocus < 0.5 μm), the number of picked particles is lower than more defocused micrographs. To test whether our workflow can pick particles on low-defocus micrographs, we selected 20 micrographs of the β-galactosidase data with estimated CTF defocus below 0.5 μm and applied the same workflow for particle picking without changing any parameters. After the iterative training, Kpicker picked 73898 particles from 213 micrographs. **Fig. 8a** is a representative micrograph with an estimated CTF defocus of 0.4 μm. Kpicker skipped the ice contaminants and picked most particles (**Fig. 8b**). Comparing to 58710 particles using only the first 20 micrographs for iterative training and picking, using low-defocus micrographs for training allowed picking 26% more particles. Therefore, our workflow might be promising for picking on low-defocus micrographs.

**Figure 8.**
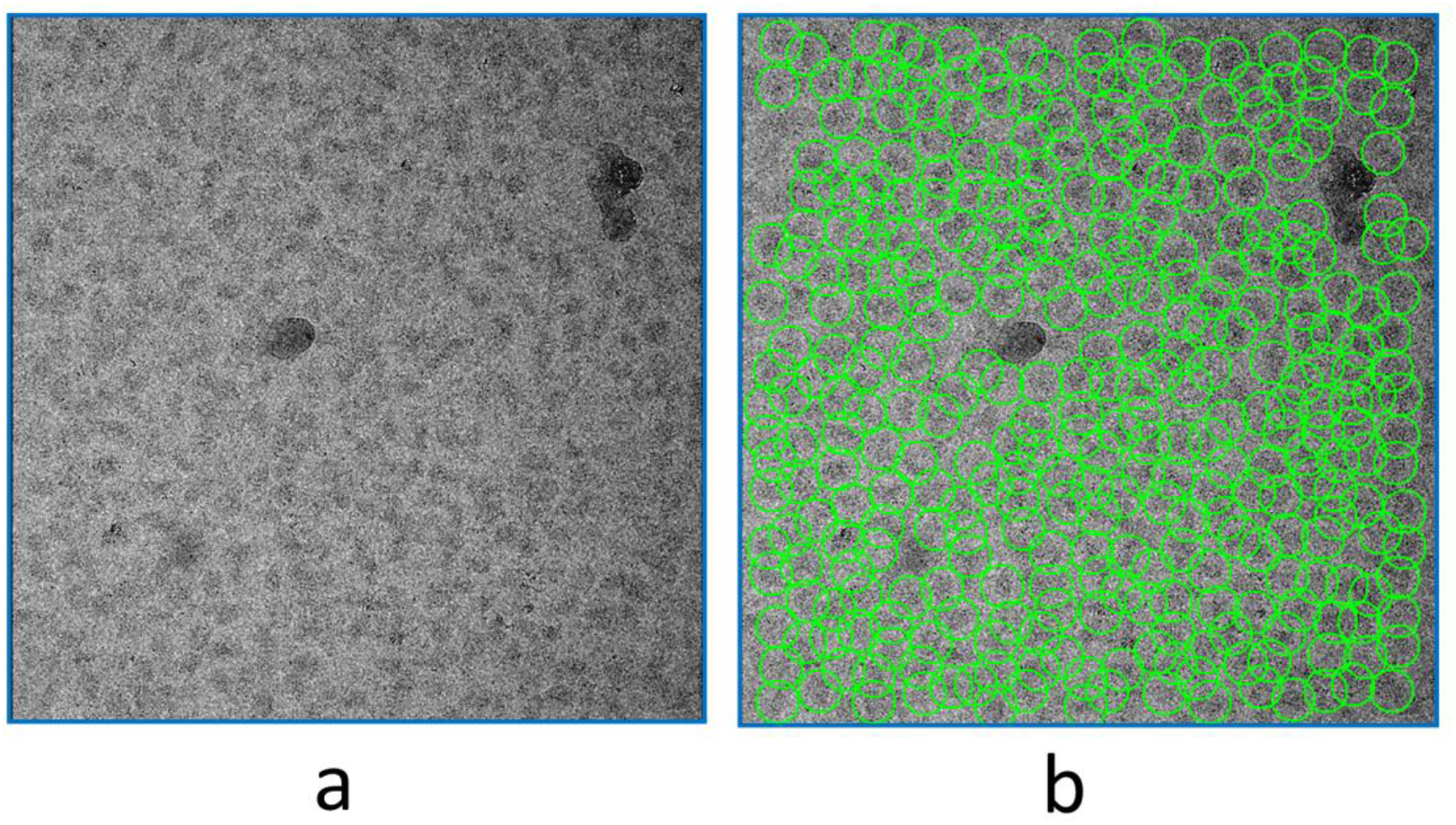
Picking on a low-defocus micrograph of the β-galactosidase data. (**a**) A micrograph with a CTF estimated defocus of 0.4 μm. (**b**) Particles picked by Kpicker after iterative training and picking with 20 micrographs of defocus values below 0.5 μm.

### 4.3 Iteration and improvement

Using 2D class average is a standard and routine procedure for cleaning up particles in single-particle Cryo-EM analysis. In our workflow, we gain two advantages from 2D class averaging. The first is to repeatedly improve training data. Such improvements may effectively remove contaminants such as ice (**Figs. 3 & 8**). The second is to use the ratio of percentage class distribution and resolution (%/Res) as a cutoff for automated selection of 2D classes. We found that %/Res is correlated well with our visual inspection and selection of 2D classes. Empirically for the six data sets, we used a value of %/Res of 0.1 for selection of 2D classes for automated iterative particle improvement and picking (**Fig. 4**). One could also use a more stringent criterion (for example %/Res > 0.2) for more difficult particle picking.

Although our workflow is devised for automated particle picking, one can also manually select 2D classes for iterative training and picking. With either automated (based on %/Res) or manual selection, improved particles may be used for Kpicker training and picking.

### 4.4 Particle picking efficiency for high-resolution reconstruction

In our workflow, we used 2D class averaging to improve particles for CNN training and picking. Therefore, we expect high percentage of picked particles will contribute to the final refinement of 3D maps. Surprisingly, for the six data sets, we found that the percentage values are quite different, from 76.5% for ribosome to only 1.8% for streptavidin (Table 1). Realizing that ribosome has a molecular weight of 1.3 MDa (Wong *et al.*, 2014) and streptavidin is one of the smallest samples tested by single-particle Cryo-EM (Han *et al.*, 2020), suggests a size dependent picking efficiency. To examine this idea more closely, we therefore plot the percentage of picked particles used for high-resolution reconstructions with respect to the molecular weight of samples used in this work (**Fig. 9**). We found there is a strong trend that the picking efficiency decreases with reduced sample molecular weight. We attribute this at least in part to beam-induced damage and denaturing at water-air interface. It is possible that smaller particles are prone to more damage and denaturing compared to large particles. Consequently, only a small portion of particles may be used for a high-resolution 3D refinement. Such damage and denaturing may not be detected at the particle picking stage that uses only low-resolution binned images. Consequently, for particles picked by the workflow, we still needed to use additional 2D and 3D classifications to filter out particles before we could reach high resolutions. Additionally, **Fig. 9** suggests that for small particles, we should expect a low picking efficiency irrespective of picking programs used.

**Figure 9.**
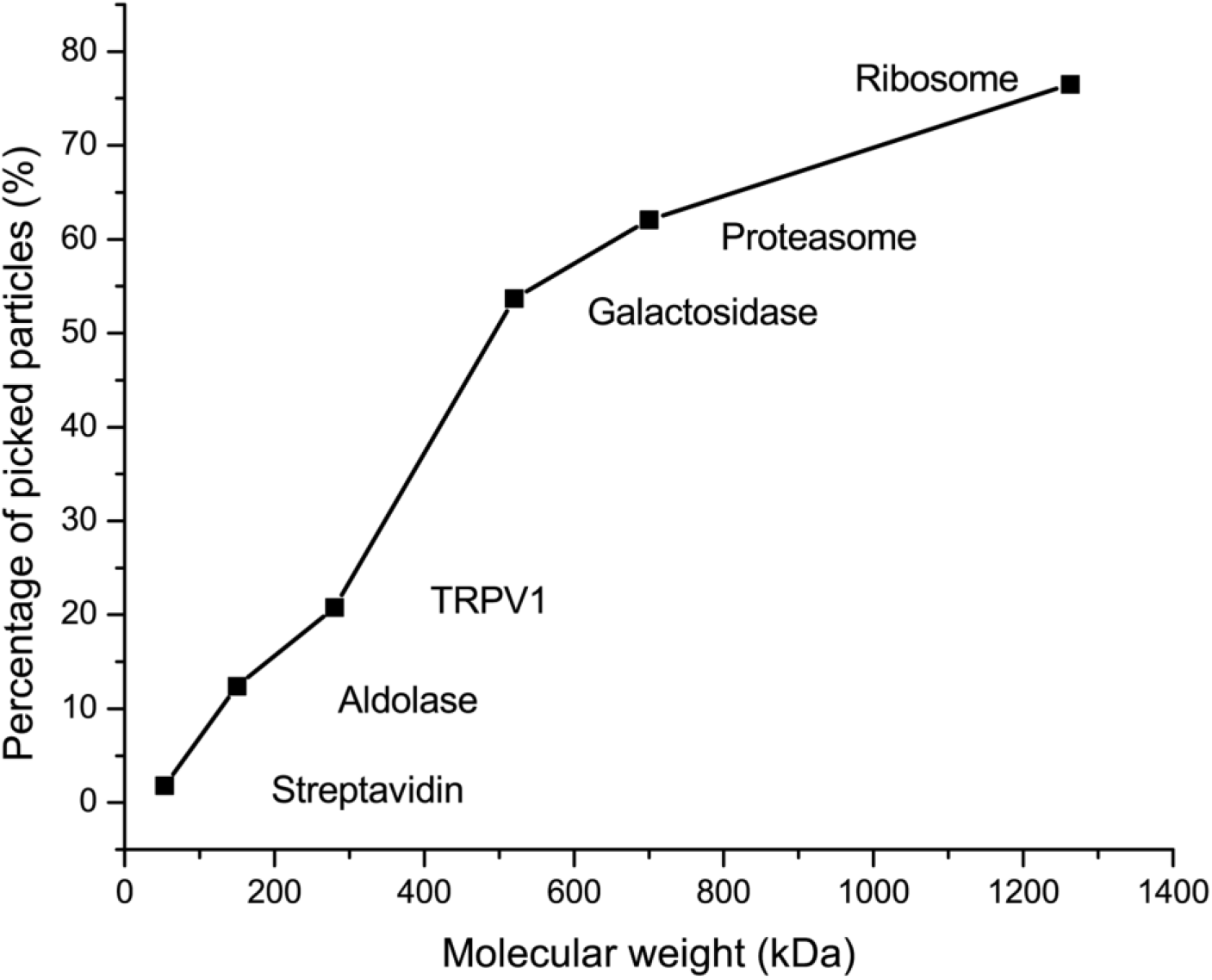
Particle-size dependent picking efficiency. The percentage of picked particles used for final map refinement was plotted with respect to sample molecular weight in kDa. For smaller-size particles, only a smaller percentage of total particles may be used for final map refinement.

One main feature of the workflow is to eliminate these trial-and-error parameters in particle picking through iteratively training a CNN model with improved particles from self-supervised 2D class average. Therefore, we did not change picking parameters in Kpicker except the particle size which is data dependent. With the six data sets, we have demonstrated that the CNN model and our workflow are highly efficient to pick sufficient quantity and quality of particles to support high-resolution reconstructions.

## 5. Concluding remarks

Particle picking is still a time-consuming step in single-particle Cryo-EM data analysis. We have proposed and tested a workflow that allows for self-supervised iterative particle picking through the integration of a deep learning-based particle picker and 2D class averaging for generation of improved training data. The workflow supports the picking of particles suitable for high-resolution single-particle analysis. Either the entire or part of the workflow may be incorporated into other workflows for automated cryo-EM single-particle analysis.

## 6. Code availability

The code of the workflow including the two pickers are available upon reasonable request.

## Acknowledgements

This work was supported by Brookhaven National Laboratory LDRD 17-023, 19-014, and the Department of Energy, Office of Science, Office of Biological and Environmental Research. D.M.M. was supported by the Virginia Ponds Scholarship Fund through the Office of Educational Program, Brookhaven National Laboratory.

